# CDK-4 regulates nucleolar size and metabolism at the cost of late-life fitness in *C. elegans*

**DOI:** 10.1101/2024.01.25.577258

**Authors:** Rachel Webster, Maria Quintana, Ran Kafri, W Brent Derry

## Abstract

An outstanding question in biology concerns mechanisms of size control in organs, cells, and organelles. Size impacts metabolic efficiency, surface area-to-volume ratio, and environmental adaptation, which are all required for optimal function. Cyclin-dependent kinase 4 (CDK4), traditionally recognized for its role in cell cycle progression, has gained increasing support for cell cycle-independent roles. Previously, we described a mechanism of cell size control involving a CDK4 and p38 MAPK circuitry that dictates target cell size. In this study, we target the CDK4/6 ortholog CDK-4 in the nematode worm *Caenorhabditis elegans* to describe functional consequences of changing biological size *in vivo*. Our data suggest that CDK-4 regulates nucleolar size and anabolic metabolism independent from cell cycle progression. When size and metabolism are increased, we report enhanced thermotolerance early in life but accelerated aging and reduced longevity late in life, suggesting a novel function of CDK-4 in somatic maintenance and organism health.

## Introduction

Cyclin-dependent kinases (CDKs), long recognized for their critical involvement in cell cycle regulation, have emerged as multifunctional enzymes boasting non-canonical roles that extend beyond the cell cycle. CDK4 is one such kinase whose role in cell cycle progression has been studied extensively. Upon activation by Cyclin D, CDK4 phosphorylates the Retinoblastoma protein (Rb) to co-ordinate the transcriptional activity of downstream E2Fs in preparation for the G1/S transition (Lees et al., 1993; Malumbres & Barbacid, 2005; Bertoli et al., 2013; Narasimha et al., 2014). As organisms age, gradual inhibition of CDK4 by p16^INK4A^ is crucial to prevent tumor growth but promotes cellular senescence due to permanent cell cycle exit (Serrano et al., 1993; Krishnamurthy et al., 2004; Rayess et al., 2012; Baker et al., 2016). New evidence of cell cycle-independent roles has begun to shift our understanding of CDK4 function, highlighting the complex interplay between the cell cycle and fundamental cellular processes, including protein synthesis, energy metabolism and stress response.

Recently, we reported that target cell size, maintained by cell size checkpoints, is specified by CDK4 activity in a dose-dependent manner. Furthermore, we showed that changes in biological size, including cell size and nucleolar size, were correlated with the activity of several metabolic pathways such as mTOR and c-Myc (Tan et al., 2021). Growing evidence has provided additional support for an active involvement of CDK4 in the regulation of cellular metabolism.

The CDK4-Rb-E2F1 axis has been implicated in the regulation of insulin secretion in pancreatic β cells and the critical balance between cell proliferation and energy metabolism (Annicotte et al., 2009; Blanchet et al., 2011). Independent from these traditional downstream targets, CDK4 has also been shown to regulate processes such as glucose production, fatty acid oxidation, and lysosomal function (Lee et al., 2014; Lopez-Mejia et al., 2017; Martínez-Carreres et al., 2019). What remains unclear from this work is the exact mechanism by which size is regulated downstream of CDK4, and whether this regulation overlaps with that of metabolism.

The appropriate production and allocation of resources is essential for proper cell division and cell growth. As organisms age, cell cycle dynamics and metabolic rate both experience significant shifts and are generally thought to intersect longevity by independent mechanisms. For example, as a central regulator of metabolism, the mTOR pathway influences the lifespan of an organism by balancing cellular processes such as protein synthesis, autophagy, and cell growth under the regulation of mTOR Complex 1 (mTORC1) (Proud, 2007; Saxton & Sabatini, 2017). Conversely, exit from the cell cycle occurs with the gradual accumulation of p16^INK4A^, inducing cellular senescence. Senescent cells promote inflammatory processes via the senescence-associated secretory phenotype (SASP), which contribute to the onset of aging and late-life pathologies (Rayess et al., 2012; Safwan-Zaiter et al., 2022). Though seemingly distinct, recent work reviewed by Wiley & Campisi, 2021 raises the idea that metabolic stress can promote senescence, leading us to hypothesize that CDK4 unites these processes under a single molecular pathway.

In the present study, we use *C. elegans* as a model to explore the influence of the CDK4/6 ortholog CDK-4 in cellular and animal physiology. The main advantages of *C. elegans* are that the CDK4-Rb-E2F axis is highly conserved, which combined with its powerful genetic tools allows for high resolution analysis of cellular effects in specific tissues. We examined nucleolar size as our primary metric for physiological size, as it directly influences translation rate and overall metabolic state of the cell. Nucleolar size changes with the rate of rRNA synthesis, which is controlled by cellular growth rate and metabolic demand (Boisvert et al., 2007; Dubois & Boisvert, 2016). Taking advantage of the Auxin-Inducible Degradation 2 (AID2) system (Negishi et al., 2022), which allows for precise spatiotemporal control over protein levels, we discovered that CDK-4 regulates nucleolar size in early larval stages independent from its role in cell cycle progression. Furthermore, we describe a novel role for CDK-4 in the regulation of protein synthesis, lipid accumulation, and stress tolerance through its substrate LIN-35 Rb, which comes at the cost of reduced longevity and healthy aging. Together, these findings represent a significant step towards a unified mechanism of aging through the co-regulation of senescence and metabolism.

## Results

### Nucleolar size increases following CDK-4 degradation, independent of cell cycle progression

Our first goal was to understand the temporal requirements for CDK-4 in the regulation of biological size *in vivo*. We previously showed that nucleolar size increased in hypodermal seam cells with the knockdown of *cdk-4* using RNA interference (RNAi). However, two generations of RNAi-mediated knockdown were required to increase size while reducing the number of cell divisions, likely owing to the role of *cdk-4* in cell cycle progression (Tan et al., 2021). We therefore asked if the nucleolar size phenotype could be recapitulated by targeting *cdk-4* within a single generation. Using the AID2 system, we targeted CDK-4 protein for conditional degradation, an advantage of this system being that proteins can be degraded rapidly with high temporal resolution during larval development (Negishi et al., 2022). Because of this, we chose to examine the seam cells given that divisions continue beyond embryogenesis and into early stages of larval development, terminally differentiating at the fourth larval stage (L4) (Sulston & Horvitz, 1977). Upon differentiation, seam cells fuse to form a syncytium that invariantly contains 16 diploid nuclei, where any deviation from this pattern of divisions strongly suggests changes in cell cycle progression. We utilized a GFP marker that localizes to seam cell nuclei and is excluded from the nucleolus (Hope, 1991), making its borders easily identifiable in the GFP-negative space. Combining this marker with the AID2 system offered the increased control over protein levels necessary to interrogate temporal requirements for CDK-4 in the regulation of nucleolar size.

Consistent with our previous report (Tan et al., 2021), CDK-4 knockdown early in larval development resulted not only in differences in nuclear count, reflecting the canonical function of CDK4 as a cell cycle regulator, but also with a significant increase in nucleolar size. When CDK-4 was ablated at the L1 stage, nucleoli were on average 1.9x larger than controls (Figure 1B). This knockdown also potently reduced the number of nuclei per seam cell syncytium, from a median of 16 in control-treated worms to 11 with the addition of auxin (Figure 1C). These influences of CDK-4 on cell count classically align with the canonical functions of CDK-4 in cell cycle progression. It also demonstrates a requirement for CDK-4 beyond early larval development for the completion of all seam cell divisions.

**Figure 1:**
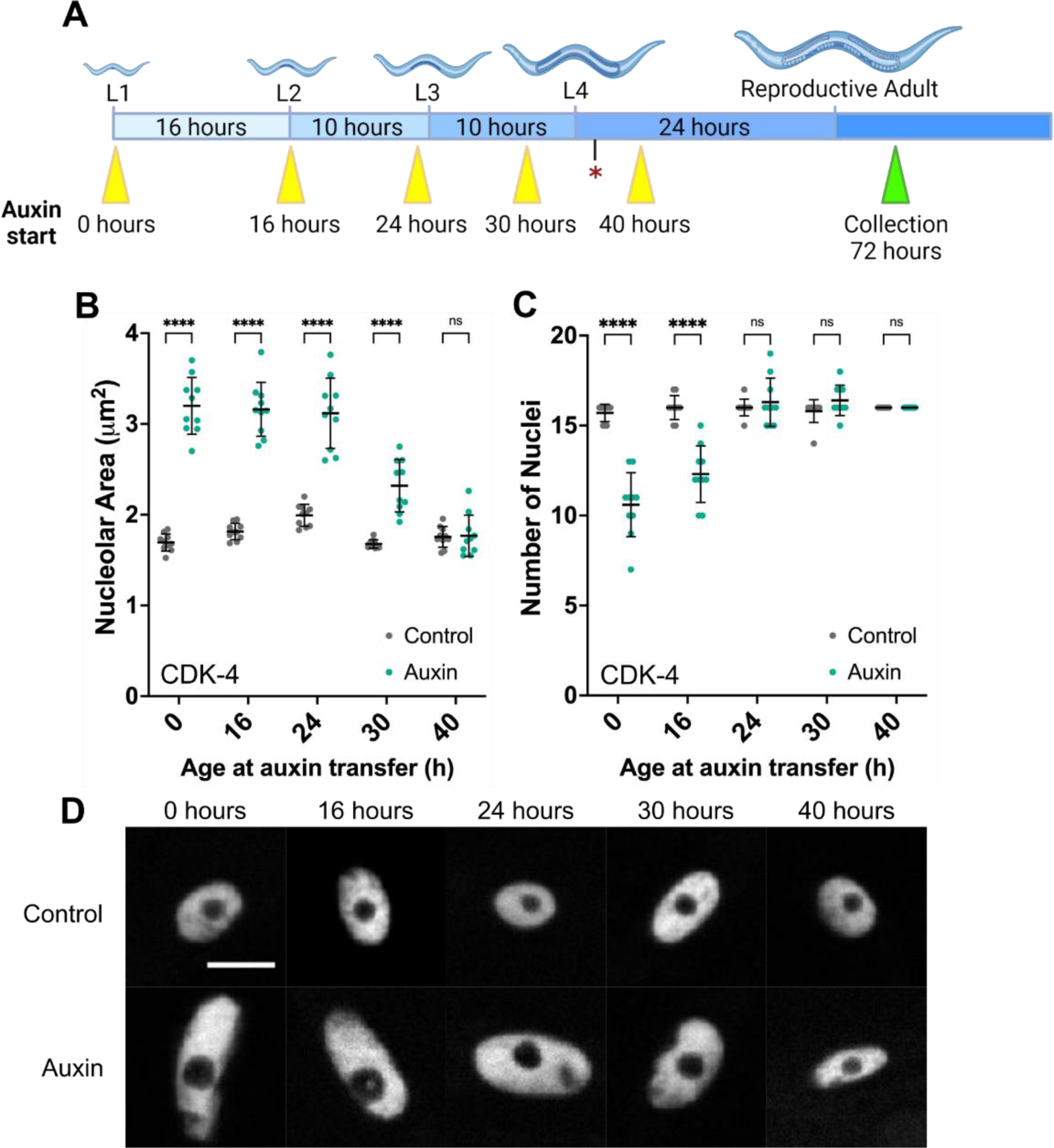
CDK-4 regulates nucleolar size with specific timing during development. **(A)** A schematic of the auxin treatment schedule followed in the time course experiment. Yellow arrows indicate the age of auxin start; time of seam cell terminal differentiation is marked with a star. Worms were grown on auxin until collection. **(B)** Nucleolar area and **(C)** number of nuclei in day 1 adults at each treatment group outlined in (A). Each point represents the average observation from all nuclei of one seam syncytium, with 10 worms measured per condition. The mean is marked with error bars showing standard deviation. Significance was calculated using two-way ANOVA with Šídák’s multiple comparisons test. **(E)** Representative images of nuclei (labelled) and nucleoli (internal circle) under each condition. Scale bar = 5µm.

Next, we asked whether the influences of CDK-4 on nucleolar size can be uncoupled from cell cycle progression. To test this, we performed a time-course experiment in which synchronized worms were transferred to auxin-supplemented plates at multiple stages of development ranging between the L1 and L4 stages, when seam cells exit the cell cycle (Figure 1A). This late-stage terminal differentiation enabled us to precisely interrogate the effects of CDK-4 ablation at different times in relation to cell cycle exit using auxin-mediated degradation. We found that nucleolar size was significantly larger than controls with CDK-4 ablation as late as 30 hours into development, with wild-type size observed at the 40-hour timepoint, corresponding with the L3 and L4 stages, respectively (Figure 1B). We also observed that nuclear size was increased in proportion to nucleolar size at each timepoint tested, suggesting that the role of CDK-4 in the regulation of biological size *in vivo* is not specific to the nucleolus (Figure S2).

Conversely, the full 16 nuclei per syncytium resulted from auxin treatment at only 24 hours of age (Figure 1C). This indicates that cell cycle progression is no longer susceptible to loss of CDK-4 after this time. Together, these experiments revealed a window of time roughly spanning the L3 larval stage when CDK-4 regulates size independent of division in the seam cells. All subsequent experiments were therefore performed with auxin treatment starting from 24 hours of age, to increase nucleolar size without affecting cell divisions. Importantly, this ability to selectively perturb size and not cell cycle progression suggests that growth and division are two distinct processes regulated by CDK-4.

### mTOR is not necessary for the increased nucleolar size observed in CDK-4 ablated worms

The mTOR pathway is a master regulator of cell growth, promoting anabolic metabolism in response to nutrient availability. To test whether the regulatory influences of CDK-4 on growth are mediated by mTORC1, we used the auxin system to deplete DAF-15, the *C. elegans* ortholog of RAPTOR. Consistent with published data, ablation of DAF-15 alone decreased nucleolar size in seam cells (Tsang et al., 2003; Tiku et al., 2017). By contrast, DAF-15 was unable to suppress the increase in nucleolar size, as the co-ablation of DAF-15 and CDK-4 resulted in enlarged nucleoli (Figure 2A). While average nucleolar size did shrink by approximately 10% in the double ablation animals, this difference fell short of statistical significance. In conclusion, our results show that DAF-15 ablation had little effect on CDK-4-mediated nucleolar enlargement, suggesting that mTORC1 and CDK-4 likely operate through parallel pathways to regulate nucleolar size.

**Figure 2:**
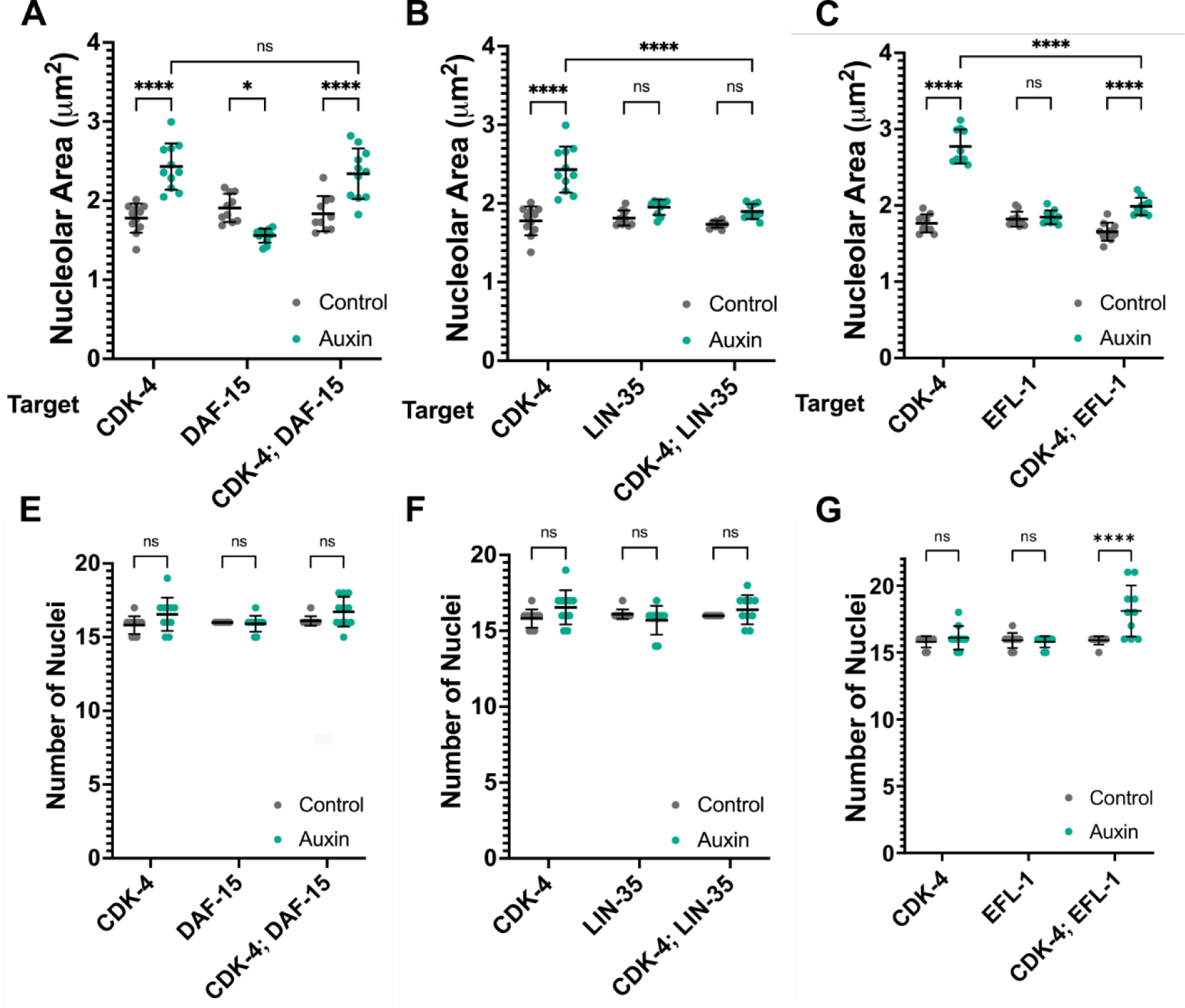
LIN-35 and EFL-1 are required for the increase in nucleolar size downstream of CDK-4. **(A)-(C)** Nucleolar area resulting from ablation of the indicated targets using auxin. Each point represents average size of all nucleoli found in one syncytium. **(D)-(F)** Number of nuclei present at the time of collection per seam syncytium. 10 worms across three biological replicates were measured per condition. Error bars denote standard deviation. Significance was calculated using two-way ANOVA with Tukey’s multiple comparisons analysis. Representative images from each condition can be found in Figure S1.

### The increase in nucleolar size caused by CDK-4 inhibition is dependent LIN-35 (Rb)/EFL-1 (E2F)

Next, we asked whether the influences of CDK-4 on nucleolar size are mediated by canonical downstream effectors LIN-35 (Rb) and EFL-1 (E2F). In the worm there is a single pocket protein, LIN-35, that most closely resembles Rb-like proteins p107 and p130 (Lu & Horvitz, 1998). To investigate the epistatic relationship between CDK-4 and LIN-35 in nucleolar size regulation, we utilized the AID2 system to target these proteins individually and jointly for degradation. We found that loss of LIN-35 alone starting from the L3 stage did not change nucleolar size and was not sufficient to inhibit seam cell divisions. Co-ablation of CDK-4 and LIN-35 similarly resulted in nucleoli of wildtype size and number, completely masking the effect of CDK-4 degradation alone (Figure 2B,E). This is consistent with previous findings showing that the CDK4 inhibitor Palbociclib increases cell size, but only in cell lines with active Rb (Ginzberg et al., 2018). These results indicate that the regulation of nucleolar size by CDK-4 is mediated by the canonical downstream target LIN-35.

To further test whether cell growth is regulated downstream of CDK-4 by the canonical CDK-4/LIN-35/EFL-1 pathway, we turned our attention to EFL-1, one of three E2Fs in *C. elegans* (Ceol & Horvitz, 2001; Winn et al., 2011). The endogenous copy of *efl-1* was tagged with the AID degron at its C-terminus, and ablation of EFL-1 starting at the L3 stage caused no changes in nucleolar size or number in the seam cells. Much like our observations with LIN-35, the effect of CDK-4 on nucleolar size was almost completely masked by the co-ablation of EFL-1 (Figure 2C). One key difference we observed under these conditions was a synthetic increase in the number of seam cell divisions among the double knockdown populations (Figure 2F). Given that size is increased even in the context of ectopic cell cycle entry, these results further support the observation that cell cycle progression and cell growth are two separate functions of CDK-4.

To ensure that the nucleolar size defect is specific to the CDK-4/LIN-35/EFL-1 pathway, and not a common feature of cell cycle inhibition, we investigated the CIP/KIP family of cyclin-dependent kinase inhibitors (CKIs) *cki-1* and *cki-2*. CKIs negatively regulate cell cycle progression with some overlap in function with LIN-35, through interactions with multiple cyclins (Sherr & Roberts, 1999; Boxem & van den Heuvel, 2001). We therefore asked if the influence of CDK-4 on nucleolar size could be suppressed by removing a negative regulator of cell cycle progression. Consistent with published data, RNAi-mediated knockdown of *cki-2* had no effect on seam cell divisions, while RNAi against *cki-1* resulted in an increase in the resulting number of nuclei (Figure S3B) (Hong et al., 1998; Boxem & van den Heuvel, 2001; Buck et al., 2009). This increase was accompanied by a modest decrease in average nucleolar size of 17% (Figure S3A). In combination with the AID2 system, knockdown of *cki-1* is not sufficient to mask the increase in size caused by loss of CDK-4 (Figure S3A). In fact, CDK-4 ablation completely prevented the excess divisions caused by *cki-1* RNAi.

Taken together, these results indicate that CDK-4 is required for maintaining physiological size of the nucleolus via the canonical LIN-35 (Rb)/EFL-1 (E2F) signaling pathway, which acts independently or downstream of TORC1. Most importantly, the effect of CDK-4 on growth is independent of its established role in cell cycle progression, revealing two independent branches of downstream signaling in this pathway.

### CDK-4 regulates protein synthesis and fat metabolism

Our results have revealed changes in nucleolar morphology after depletion of CDK-4. Nucleolar size correlates with the synthesis of ribosomal RNAs, and thus the production of new ribosomes (Ma et al., 2016). We therefore hypothesized that the synthesis of new proteins, and perhaps additional aspects of cellular metabolism, are consequently upregulated when CDK-4 is lost. To explore this possibility, we investigated several biosynthetic processes. Turning first to protein synthesis, we employed the SUnSET (SUrface SEnsing of Translation) assay, which uses Puromycin incorporation into nascent peptide chains as a readout of mRNA translation during a defined period of time. We found that ablation of CDK-4 resulted in puromycin incorporation at a rate of about 80% higher than controls when normalized to total protein (Figure 3B). Furthermore, the increase in puromycin incorporation appears to be uniform across all proteins, suggesting that this increase is occurring globally across the proteome (Figure 3A).

**Figure 3:**
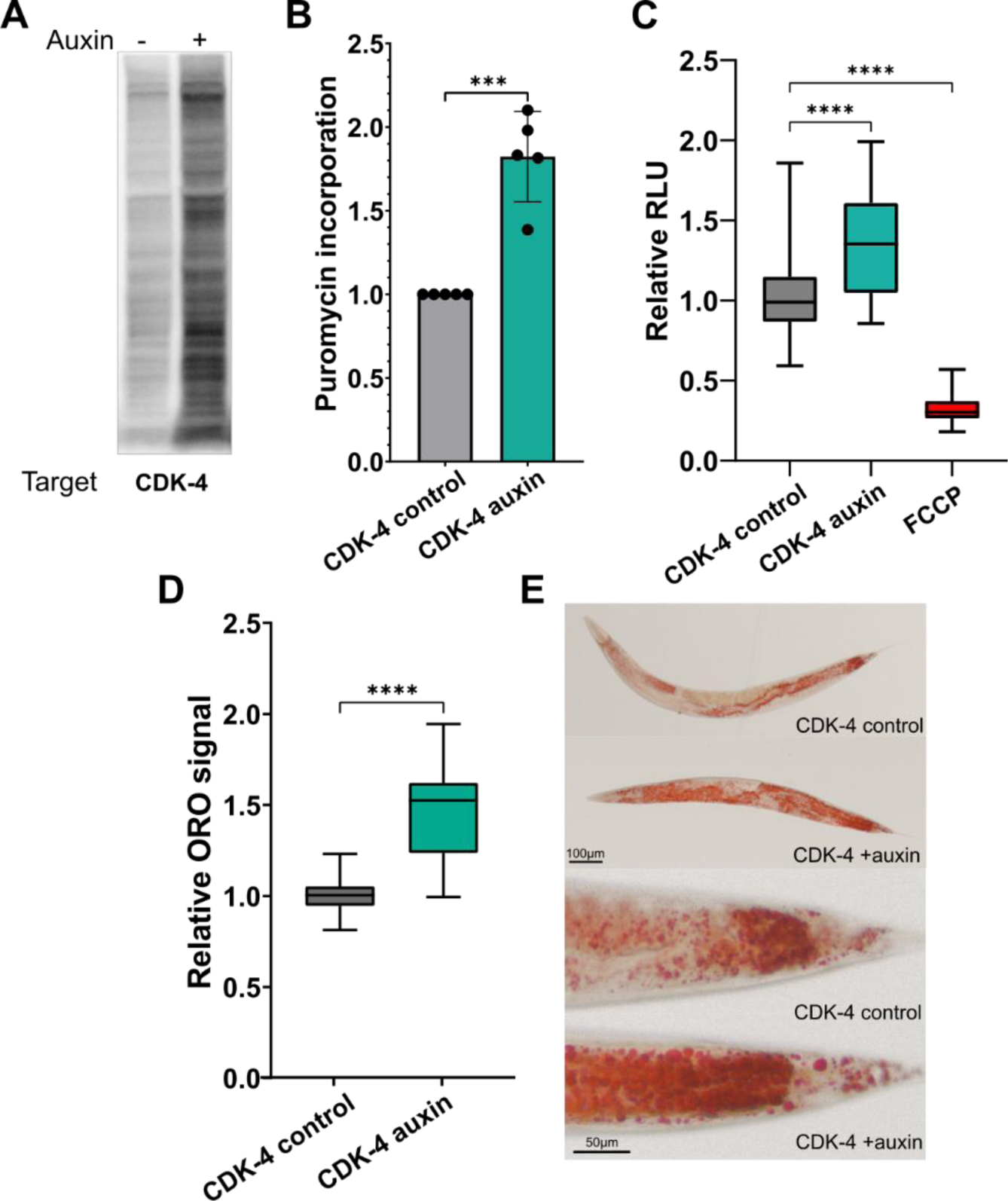
CDK-4 regulates anabolic metabolism. **(A)** Representative western blot showing puromycin incorporation estimating translation rate under control or auxin treatment. SYPRO Ruby loading control in Figure S4A. **(B)** Quantification of five biological replicates of the sunset assay normalized to total protein and visualized relative to control. Bar height represents the mean with error bars denoting standard deviation. Significance was calculated using Student’s t-test, p = 0.00013. **(C)** Quantification of luminescence (relative light units) as an estimation of ATP levels between auxin and control populations, using FCCP treatment as a positive control. Three biological replicates are represented normalized to control, with error bars showing minimum and maximum values. Significance was calculated using one-way ANOVA with Dunnett’s multiple comparisons analysis. **(D)** Quantification of Oil Red O staining across three biological replicates. Error bars capture minimum and maximum values. Significance was calculated using Student’s t-test. **(E)** Representative images of whole worm ORO staining on top and staining in tail regions below.

We next asked if ATP levels were elevated to support increased protein synthesis using a luciferase-based method developed to assess steady state ATP levels in live worms (Lagido et al., 2008). These worms express firefly luciferase ubiquitously in the cytoplasm, and luciferin salt is consumed with an *E.coli* food source mixed into the media. Since ATP is the limiting reagent in this reaction, the amount of measured luminescence is proportional to levels of intracellular ATP. Figure 3C indicates that CDK-4 ablation resulted in a considerable increase in luminescence over controls. This shows that the amount of ATP produced in CDK-4-ablated worms is greater and implies a shift in energy homeostasis resulting from the loss of CDK-4.

Since protein biosynthesis and ATP levels were upregulated in the absence of CDK-4, we wondered if lipid production would follow the same trend. Using Oil Red O, we stained whole worms for neutral lipid content following CDK-4 ablation. With this method, we observed significantly increased fat deposits across the whole worm by day 1 of adulthood, with large droplets accumulating in the body cavity, and enrichment also detectable in the intestine (Figure 3D,E). Taken together, these results show that the increased synthesis of proteins and lipids suggests that CDK-4 normally acts to limit anabolic processes in order to maintain proper cellular function.

### Regulation of anabolic metabolism by CDK-4 requires LIN-35/Rb, but not mTORC1

Our next goal was to determine whether the metabolic phenotypes resulting from CDK-4 knockdown relied on the same downstream mechanism as size regulation. Figure 2A presents compelling evidence that the mTORC1 pathway is not a prerequisite for the large nucleolar size resulting from CDK-4 ablation. This led us to ask whether CDK-4’s influence on protein synthesis similarly bypasses mTORC1 dependency. Employing the SUnSET assay, we evaluated protein synthesis in worms devoid of both CDK-4 and DAF-15. Aligning with the known functions of mTORC1, ablation of DAF-15 produced a 25% decrease in puromycin signal intensity (Figure S5A,B). By contrast, the co-ablation of CDK-4 and DAF-15 resulted in an approximately 40% increase in translation, and as such was not capable of significantly suppressing the CDK-4-dependent translation phenotype. This result indicates that DAF-15 acts independently of CDK-4 to regulate mRNA translation.

In addition to regulating nucleolar size and protein synthesis, mTORC1 is also a key regulator of lipid synthesis via SREBP1 (Porstmann et al., 2008). We therefore investigated whether lipid accumulation following CDK-4 ablation might also be dependent on DAF-15. Our experiments revealed that the combined ablation of DAF-15 with CDK-4 did not prevent the rise in lipid accumulation, suggesting that these metabolic processes are regulated downstream of CDK-4 and mTORC1, but through parallel pathways (Figure S5C). These findings are consistent with the observed changes in nucleolar size and further substantiate the notion that CDK-4 independently influences anabolic metabolism, separate from mTORC1. Additionally, these results not only reinforce but also expand our understanding of the anabolic functions of CDK-4 as being distinct from the mTORC1 pathway.

We next investigated whether LIN-35 Rb was required for the metabolic effects of CDK-4. LIN-35 has not been previously linked to metabolic regulation in the worm, however some studies have implicated the orthologous Rb/E2F axis in maintaining energy homeostasis in other systems (Blanchet et al., 2011; Denechaud et al., 2016; Mandigo et al., 2021). Consistent with nucleolar size, we found that ablation of LIN-35 alone did not affect translation. However, its combined ablation with CDK-4 completely suppressed the increased translation caused by loss of CDK-4 alone (Figure 4A,B). The same trend was observed for both ATP levels, for which LIN-35 ablation on its own had no effect but was able to mask the phenotype of CDK-4 (Figure 4C). Loss of LIN-35 showed a modest increase in lipid levels, however its co-ablation with CDK-4 significantly suppressed the increase in fat accumulation caused by CDK-4 ablation alone (Figure 4D,E). Overall, LIN-35 is required for mediating all tested metabolic processes regulated by CDK-4.

**Figure 4:**
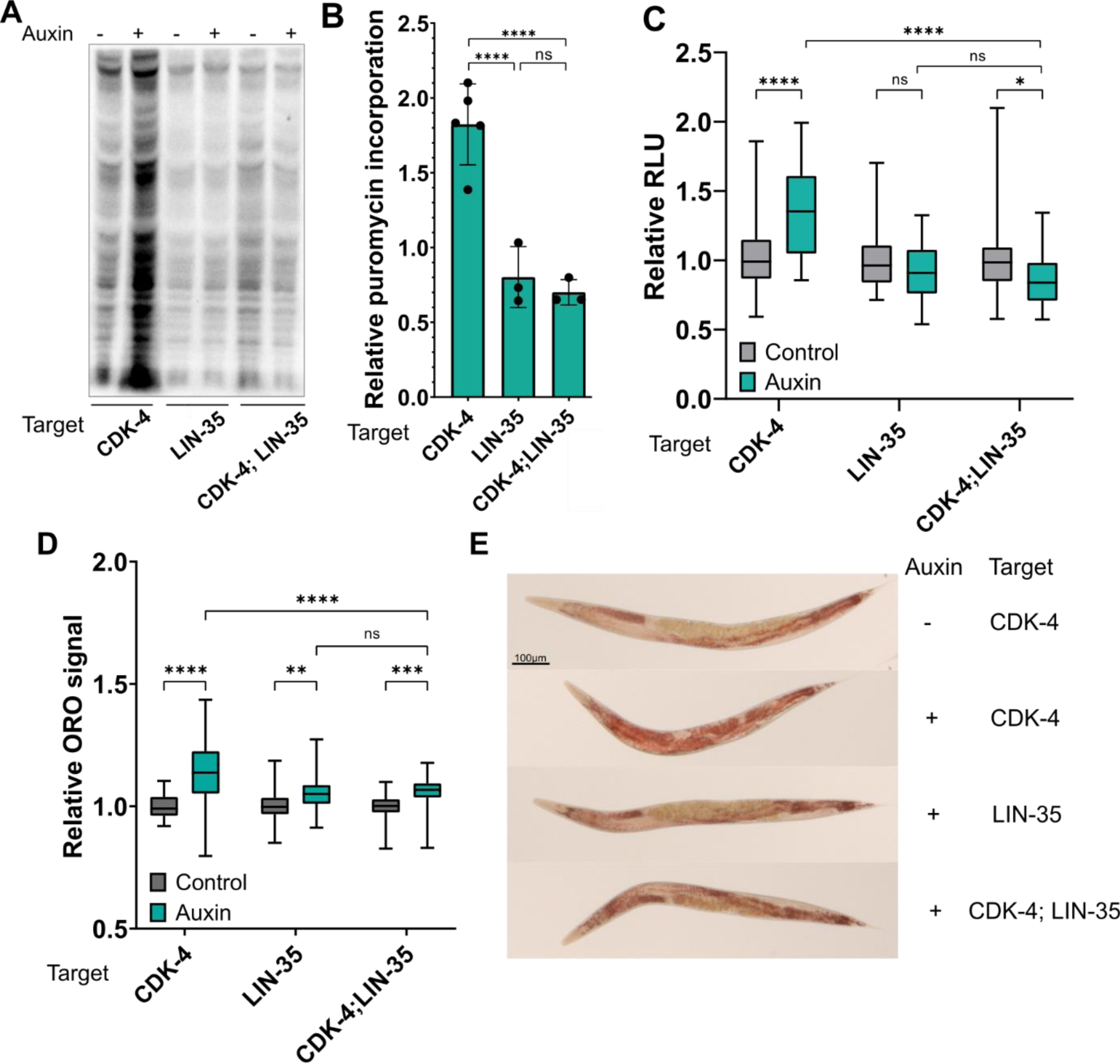
The effect of CDK-4 on biosynthesis requires LIN-35. **(A)** Representative western blot showing puromycin incorporation across the indicated conditions. SYPRO Ruby loading control in Figure S4B. **(B)** Quantification of at least three biological replicates of the sunset assay normalized to total protein and visualized as fold-increase over controls. Significance was calculated using one-way ANOVA with Dunnett’s multiple comparisons analysis. **(C)** Quantification of luminescence (relative light units) to measure ATP levels between auxin and control populations of the indicated targets. Three biological replicates are represented normalized to control, with error bars representing minimum to maximum values. Significance was calculated using two-way ANOVA with Tukey’ multiple comparisons analysis. **(D)** Quantification of Oil Red O staining in between 49-67 worms per condition across three biological replicates of the indicated conditions. Error bars denote the minimum to maximum values. Significance was calculated using two-way ANOVA with Tukey’s multiple comparisons analysis. **(E)** Representative images of whole worm ORO staining.

### CDK-4 ablation increases tolerance of temperature stress

The nucleolus is a multifunctional organelle, with ribosome subunit assembly being only one of its primary functions. It is estimated that almost 70% of nucleolar proteins are involved in processes unrelated to ribosome biogenesis, such as mRNA processing, DNA repair, cell cycle control, and cellular stress response (Andersen et al., 2005). Changes in nucleolar size and morphology are often observed in response to stress (Boisvert et al., 2007; Boulon et al., 2010; Frottin et al., 2019), so we next asked whether the ability to survive and recover from acute stress changes with increased nucleolar size following CDK-4 ablation.

Worms were exposed to elevated temperatures for a duration of time sufficient to cause lethality in wildtype animals. L4-stage worms were shifted from 20°C to 37°C for four hours, allowed to recover overnight, and survival quantified. In control populations under these conditions, up to 10% of the population will survive this heat shock. However, after CDK-4 ablation 56% of the population survived the same heat shock treatment (Figure 5B). Although LIN-35 ablation had no significant effect on heat stress survival, co-ablation with CDK-4 suppressed survival to wildtype levels (Figure 5B). These results suggest that when nucleoli are large and protein synthesis is high, the animal is better equipped to survive phases of acute heat stress.

**Figure 5:**
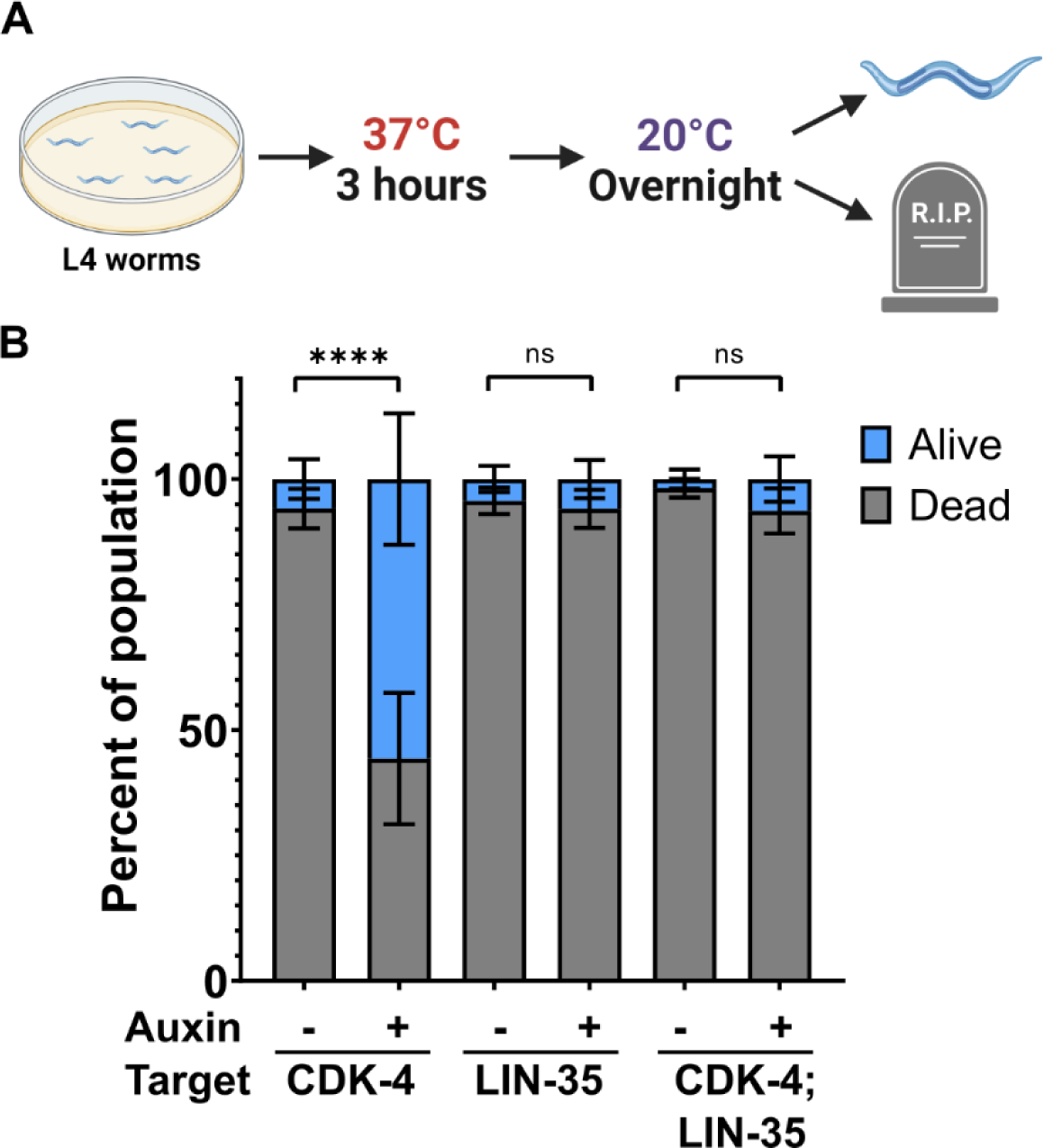
CDK-4 ablation leads to increased heat stress survival early in life. **(A)** Schematic outlining the process of heat shock treatment. **(B)** Proportions of the indicated populations that survived extreme heat stress. Three biological replicates using populations of 50 worms each are represented, with bar heights representing the mean and error bars denoting standard deviation. Significance was calculated with two-way ANOVA using Šídák’s multiple comparisons test.

### Loss of CDK-4 accelerates aging

Protein synthesis has long been associated with longevity (Hansen et al., 2007; Pan et al., 2007; Martinez-Miguel et al., 2021; Kim & Pickering, 2023), but recent work has described nucleolar size regulation as a converging point for diverse longevity mechanisms. Support for this comes from the observation that longevity is inversely correlated with nucleolar size, where small nucleoli are generally correlated with increased lifespan in *C. elegans* (Tiku et al., 2017). Furthermore, premature aging in Hutchinson-Gilford progeria syndrome (HGPS) patients was found to be associated with large nucleoli and elevated protein synthesis (Buchwalter & Hetzer, 2017). Independently, CDK4 has been implicated in the rate of aging through the onset of senescence. Inhibition of CDK4 activity in mammals by p16^INK4A^ causes irreversible cell cycle arrest, morphological changes, and the secretion of pro-inflammatory factors, all hallmarks of cellular senescence that shorten lifespan (Rayess et al., 2012; Baker et al., 2016). Given that we have defined the additional role for CDK-4 in the regulation of nucleolar morphology and function under conditions that do not impede cell divisions, we wondered whether these phenotypes were negatively affecting survival and late-life health of CDK-4-ablated worms.

CDK-4 ablation decreased median lifespan (Figure 6A), but on its own, reduced lifespan is not sufficient evidence for accelerated aging. In addition to lifespan, we therefore examined several established readouts of physical degradation and aging in young adult populations as a proxy for how well worms endure with age. First, we quantified movement in liquid and rate of pharyngeal contractions, which decline with muscle degeneration in aging *C. elegans* (Herndon et al., 2002; Fisher, 2004; Huang et al., 2004). By day one of adulthood, CDK-4 ablated populations exhibited wildtype movement, however by day three there was a 50% reduction in body bends per minute, progressively declining with time at a rate faster than controls (Figure 6B). We also observed normal pharyngeal contraction rate in early adulthood, with a sharp decline by day three (Figure 6C). These processes naturally decline with age, however with loss of CDK-4 we observed premature degeneration associated with the reduction in lifespan.

**Figure 6:**
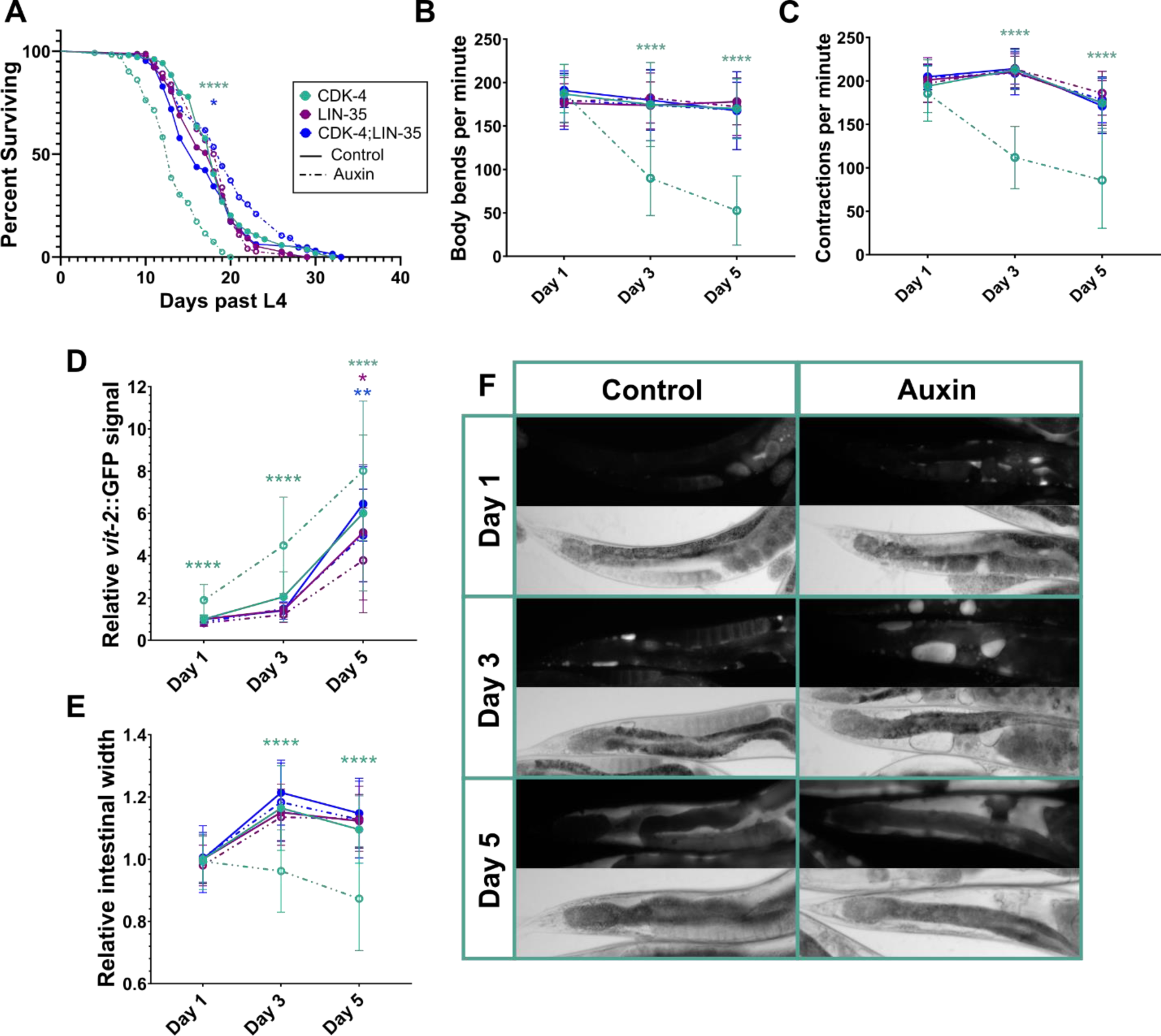
Loss of CDK-4 decreases longevity and accelerates rate of aging and degeneration. **(A)** Survival of strains with the listed proteins (box) targeted by AID2. Data shown from a minimum of 64 worms per condition across at least two biological replicates. Significance was calculated using the log-rank (Mantel-Cox) test, *p = 0.0299. **(B)** Thrashing rate in liquid and **(C)** rate of pharyngeal contractions in worms aged to the indicated time points. Significance from at least 15 worms across three biological replicates was calculated using two-way ANOVA with Tukey’s multiple comparisons analysis. **(D)** Quantification of total VIT-2::GFP accumulation and **(E)** posterior intestinal width across three biological replicates and normalized to day 1 controls for each condition, with 39-64 worms measured per condition. Significance was calculated using two-way ANOVA with Tukey’s multiple comparisons analysis, *p = 0.0153, **p = 0.0022. Unlabelled p-values are not significant. **(F)** Representative images of VIT-2::GFP accumulation (top) and intestinal width (bottom) at each time point in *cdk-4*::degron-expressing worms.

To understand if premature aging was detectable in other tissues, we quantified the accumulation of yolk protein VIT-2. Increased yolk accumulation is a hallmark feature of the normal aging process in *C. elegans* and correlates with the decline in fecundity during mid-adulthood (Ezcurra et al., 2018). By day five of adulthood, control worms on average accumulated 2.6x more yolk protein compared to day one adults. In CDK-4 depleted animals, yolk accumulated at an accelerated rate, with more than a 2-fold increase over controls in day one adults (Figure 6D,F). Yolk secretion originates from intestinal cells and is accompanied by a decrease in intestinal width as yolk is produced, which we found to decline earlier in CDK-4 ablated worms than in age-matched controls (Figure 6E). Together, these results indicate an early decline in healthspan and decreased lifespan in worms lacking CDK-4.

Since LIN-35 is required for all phenotypes induced by loss of CDK-4, including nucleolar size, metabolism, and stress resistance, we asked if the reduced longevity and accelerated aging of CDK-4 ablated worms was dependent on LIN-35. Although loss of LIN-35 alone had no significant effect on lifespan or healthspan, its co-ablation with CDK-4 was able to rescue lifespan and all hallmarks of aging (Figure 6A-E). Collectively, these results indicate a cell cycle independent role for CDK-4 in protein and lipid biosynthesis that impacts lifespan and healthspan through its canonical effector LIN-35/Rb.

## Discussion

CDK4 has been predominantly studied for its role as a cell cycle regulator, where it is required for promoting the transition into S phase through direct phosphorylation of Rb. We report a novel role for CDK-4 in the regulation of nucleolar size and anabolic metabolism, in a way that can be dissociated from cell cycle progression. Our data demonstrates that CDK-4 functions to limit the synthesis of proteins and lipids during development via canonical effectors LIN-35 (Rb)/EFL-1 (E2F). Recent observations support the idea of cell cycle-independent functions of CDK4 in other models, suggesting that CDK4 functions as a conserved regulator of systemic metabolism. For example, CDK4 can influence AMPK, PGC-1α and mTORC1 activity through Rb-independent interactions, regulating cellular processes such as glucose production and lysosomal function (Lees et al., 1993; Martínez-Carreres et al., 2019; Romero-Pozuelo et al., 2020). Other roles beyond cell cycle progression have been shown to be mediated by canonical effectors Rb and the E2Fs. Insulin secretion and oxidative metabolism was shown to be upregulated upon loss of E2F1, in a way that could be rescued by CDK4 gain-of-function (Blanchet et al., 2011). In pancreatic β-cells that are not actively dividing, insulin secretion is promoted by CDK4-Rb-E2F activity through transcriptional regulation of glucose regulator Kir6.2 (Annicotte et al., 2009). However, CDK4 inhibition by p16^INK4A^ also promotes insulin production involving mTOR and PPAR-γ (Helman et al., 2016), suggesting that the dynamics between CDK4 activity and energy metabolism are highly nuanced and perhaps context-dependent.

Here, we have shown that CDK-4 regulates nucleolar size and anabolic metabolism in *C. elegans* without impacting the number of cell divisions. Through its canonical effectors LIN-35 (Rb)/EFL-1 (E2F), CDK-4 functions to limit the synthesis of proteins and lipids during development. Ablation of CDK-4 resulted in enlarged nucleoli and increased metabolism, which we found reduces lifespan and accelerates aging. One question that has remained unclear is whether the regulation of nucleolar size and metabolic processes are secondary to the cell cycle defects, or if they represent strictly independent functions for CDK4. Importantly, we demonstrate that changes in size caused by the ablation of CDK-4 are apparent when cell cycle progression is unperturbed. Ablation of CDK-4 during the mid-larval stages of development revealed a clear uncoupling of nucleolar size regulation from cell cycle progression. However, we also found that ablating CDK-4 starting at 40 hours of age had no effect on nucleolar size. The timing of this coincides with exit from the cell cycle in seam cells at the L4 stage (Sulston & Horvitz, 1977). These specific temporal effects on size therefore led us to conclude that although size and cell cycle progression appear to be independently regulated downstream of CDK-4, it is likely that size is mediated by canonical cell cycle machinery.

An advantage of using *C. elegans* as a model for this study is the availability of genetic tools for conducting epistasis experiments, which allowed us to test the requirement of known CDK-4 targets and downstream effectors at specific times during development. Given the link between CDK4 and mTOR described in previous studies, and the role of mTOR in regulating both metabolism and nucleolar dynamics (Boulon et al., 2010; Blackwell et al., 2019; Martínez-Carreres et al., 2019; Romero-Pozuelo et al., 2020), we were surprised to find that CDK-4 regulates nucleolar size and metabolism independently of mTORC1. We hypothesized that upregulation of mTORC1 activity was responsible for these phenotypes, but ablation of DAF-15 did not suppress nucleolar size and growth phenotypes caused by loss of CDK-4. This suggests that mTORC1 regulates these same metabolic processes in parallel to CDK-4.

Contrary to expectations, our results indicate that nucleolar size and the subsequent biosynthetic phenotypes in *C. elegans* are entirely dependent on the CDK-4 substrate LIN-35/Rb. This observation agrees with previous data in mammalian cells, demonstrating that the effect of CDK4 on size requires Rb (Ginzberg et al., 2018). However, despite being regulated by canonical cell cycle machinery, the size phenotype appears to be specific to CDK4/CDK-4 rather than a general consequence of perturbing cell cycle progression. Although CDK2 and CDK4 both phosphorylate Rb to promote the G1/S transition, and inhibition of either CDK is sufficient to lengthen cell cycle duration, only the inhibition of CDK4 caused an increase in cell size (Tan et al., 2021). Furthermore, we showed here that loss of the cyclin-dependent kinase inhibitor *cki-1* could not rescue the nucleolar size phenotype caused by CDK-4 ablation, providing further support that these phenotypes are independent of cell cycle progression.

Loss of EFL-1 together with CDK-4 was able to significantly suppress the increase in nucleolar size, albeit not entirely to control levels, as seen with LIN-35 co-ablation. This might be explained by the existence of two additional E2F-like proteins in the worm, EFL-2 and EFL-3 (Ceol & Horvitz, 2001; Winn et al., 2011). While loss-of-function mutations in either *efl-1* or *efl-2* are viable, *efl-1;efl-2* double mutants are lethal, indicating redundant roles in an essential process (Schertel & Conradt, 2007). EFL-3 is described as an ortholog of E2F7 and E2F8, atypical in that they do not associate with Rb (Winn et al., 2011), therefore it is an unlikely candidate for mediating these phenotypes. However, it is possible that EFL-2 provides some level of compensation when EFL-1 is lost. In addition, human Rb exists in complex with E2F proteins as well as more than 100 additional binding targets (Morris & Dyson, 2001). Interactions between LIN-35 and any additional transcription factors independent from the E2Fs might explain how EFL-1 ablation alone cannot fully rescue the nucleolar size phenotype.

Protein synthesis, lipid biosynthesis, aging, and stress phenotypes are likely linked to the observed changes in nucleolar size and function. Large nucleoli produce more ribosomes, increasing the capacity of a cell to translate the proteins necessary for cellular functions during development and in response to stress. Nucleolar morphology and protein synthesis have been specifically linked with aging and lifespan, as small nucleoli and conditions that restrict mRNA translation are characteristic of long-lived individuals (Hansen et al., 2007; Tiku et al., 2017; Martinez-Miguel et al., 2021; Kim & Pickering, 2023). The relation between lipid levels and lifespan is not so straightforward among long-lived mutants, with *daf-*2/IGF1R and *glp-1* germline-defective mutants exhibiting increased lipid stores, while *eat-2* caloric restriction mutants have reduced fat levels (Kimura et al., 1997; Lakowski & Hekimi, 1998; O’Rourke et al., 2009). With the ablation of CDK-4 we see the expected increase in protein synthesis given large nucleolar size and reduced lifespan, which agrees with previous reports. This perturbation also causes an increase in lipid accumulation, demonstrating a broad increase in biosynthesis, however a direct link between this accumulation and the observed aging phenotypes has yet to be determined.

Although the ablation of CDK-4 resulted in accelerated aging, these worms demonstrated an increased thermotolerance early in life. Our work suggests that an increase in nucleolar size and perhaps biosynthesis act protectively early in life, but significantly reduce fitness and survival with age. This trade-off was unexpected, given that long-lived mutants with reduced mTOR or insulin signalling, associated with decreased nucleolar size and protein translation, exhibit a broad resistance to stress (Hansen et al., 2007; Zhou et al., 2011; Tiku et al., 2017).

While the exact cause of this protection against acute heat stress is unknown, we hypothesize that it represents a novel benefit of large nucleoli not before shown *in vivo*. A possible explanation for this concerns the nucleolus, recently reported to act protectively in times of heat stress by sequestering misfolded proteins to prevent aggregate formation and allow for efficient refolding during recovery (Frottin et al., 2019). Perhaps large nucleoli, consequential of CDK-4 ablation, have an increased capacity to harbour misfolded proteins and increase chances of survival following acute heat stress.

We have shown that CDK-4 ablation causes not only a reduction in lifespan, but an accelerated onset of aging phenotypes. Centerstage to mechanisms of aging are two distinct molecular processes: metabolic rate and cellular senescence. The major metabolic pathways such as mTOR, insulin signaling and c-Myc are all well-established regulators of longevity, where decreased activity in any of these pathways significantly extends lifespan (Hofmann et al., 2015; Dall & Færgeman, 2019; Papadopoli et al., 2019). Conversely, cellular senescence is a hallmark of aging, promoting morphological and chemical changes that contribute to the aging process (Kumari & Jat, 2021). CDK4 is classically involved in senescence through its direct inhibition by p16^INK4A^. Increasing levels of p16 are found in aging animals, while specific removal of p16 is sufficient to increase lifespan and healthspan (Rayess et al., 2012; Baker et al., 2016). Here, we showed that loss of CDK-4 in worms accelerates aging and shortens lifespan consistent with mammalian evidence, however we also identified a direct link between CDK-4 and metabolic rate. While senescence and metabolism have traditionally been considered independent influences on lifespan, the common involvement of CDK-4 in both branches suggests that these processes are commonly regulated through a single molecular pathway.

Collectively, our results support the idea that nucleolar size and metabolism are strongly dependent on the canonical G1/S signaling axis of CDK-4/LIN-35/EFL-1, but independent of cell cycle progression. This work provides new insights into how CDK-4 and its canonical signaling pathway coordinate cell cycle and metabolism to optimize developmental robustness and organism lifespan.

## Acknowledgements

This work was supported by a Canadian Institutes of Health Research Project grant to WBD. (PJT 165837). WBD is the Canada Research Chair in Animal Models of Human Disease.

## Methods

### Strains and maintenance

Nematode strains were maintained at 20°C on nematode growth media (NGM) agar plates seeded with OP50 *E. coli* bacteria unless otherwise stated. Some strains were provided by the CGC, which is funded by NIH Office of Research Infrastructure Programs (P40 OD010440). DV3525 [*daf-15(re257[daf-15::mNG::AID])* was a kind gift from the Reiner lab (Duong et al., 2020).

### CRISPR-Cas9 mutagenesis

Mutations generated using CRISPR mutagenesis were created following the single-stranded donor oligonucleotide method described in Eroglu *et al*. (Eroglu et al., 2023). Mutations were generated in the HS3545 background, which ubiquitously expresses the TIR1(F79G) receptor, as well as *elt-3*p::GFP. Single-stranded donor oligonucleotides were either ordered as such for shorter insertions or created from a double-stranded template for longer insertions. In the case of longer insertions, one of two amplifying primers was phosphorylated on the 5’ end. Following amplification, the resulting product was digested by exonuclease to obtain the single-stranded template. Day 1 adult worms were microinjected with a mix containing sgRNA-loaded cas9 protein, single-stranded donor oligonucleotide, and sgRNA and repair template targeting *dpy-10* used as a co-injection marker for screening progeny. Edits were confirmed by sequencing.

### Auxin treatment

Populations of worms were synchronized by dissolving gravid adults in a hypochlorite solution (20% v/v sodium hypochlorite, 25% v/v sodium hydroxide, and 55% v/v distilled water), washing 3-4x in M9 buffer, and allowing L1s to hatch overnight in fresh M9. L1s were then plated on NGM plates for the time described in each experiment, then divided between control and auxin plates. To make auxin plates, NGM agar media was supplemented with 2.5μM 5-Ph-IAA (MedChemExpress 168649-23-8) or 0.3% Dimethyl sulfoxide (DMSO) for controls. Aging experiments that required maintaining worms through the egg-laying phase were further supplemented with 7.5μM C22 (Chembridge 9345554) for both control and auxin-treated conditions. C22 disrupts egg viability without harming adult worms (Weicksel et al, 2016). Plates were seeded using 10x OP50 grown in LB overnight at 37°C, 80μL on small plates or 250μL on medium plates.

### RNA interference

Worms were fed dsRNA-expressing HT115 *E. coli* bacteria to knock down targets of interest. Strains were grown from the Source Bioscience LifeSciences feeding library. RNAi plates (NGM supplemented with 25μg/mL carbenicillin and 2.5mM IPTG) were seeded with induced RNAi bacteria. 50 L1 worms per condition were plated and grown to the adult stage. 10-15 gravid adults were transferred to fresh RNAi plates for a timed egg laying of 2 hours to synchronize progeny before being removed. Progeny were transferred to auxin or control plates (supplemented with 25μg/mL carbenicillin and 2.5mM IPTG) at 24 hours post-hatching and collected for nucleolar size measurements at 72 hours.

### Nucleolar size measurement

The SCMp::GFP marker from strain JR667 was crossed into the relevant AID strains to mark seam cell nuclei. Strains were bleached to synchronize and treated with auxin, transferred from NGM at 24 hours past L1. At 72 hours, worms were mounted on 4% agarose slides in 100mM tetramisole. Using a Zeiss Axio Observer.Z1 microscope, worms were imaged in the eGFP channel at 63x, using z-stacks to capture the midpoint of each nucleolus in the field of view. Stacks were exported using Zen Pro imaging software. The layer capturing the maximum cross-sectional area was manually traced and measured using ImageJ.

### SUnSET assay

Protocol adapted from (Arnold et al., 2014). 40,000 synchronized worms were obtained by bleaching (as described above) four medium NGM plates seeded with OP50. L1s were plated on one large NGM plate seeded with 1mL 30x OP50 grown in Terrific Broth (TB). After 24 hours, L3 larvae were washed from the plates using M9. The collected worms were transferred between a large auxin and large control plate seeded with 1mL concentrated OP50, aiming for 16,000 worms per condition. At 72 hours, worms were washed off plates using S Basal and collected in 15 mL Falcon tubes, where they were washed 3x in S Basal to remove excess bacteria. Worms were resuspended with 600µL 10x OP50, 180µL of 10mg/mL puromycin stock solution (Thermofisher A1113803) and S Medium up to 4.5mL. The final concentration of puromycin was 0.4mg/mL. Samples were rocked at room temperature for 4 hours. Following this pulse, worms were washed 3x with S Basal, and a final wash in ddH2O, removing as much supernatant as possible to prepare for protein collection.

Protein was collected by lysing worms in a 1:1 ratio of worm pellet to lysis buffer using a tissue grinder (VWR 62400-675) kept cold on ice. Lysis buffer contained 30mM HEPES pH 7.5, 100mM Potassium acetate, 2mM Magnesium acetate, 1 tablet/5mL cOmplete^TM^ mini EDTA-free protease inhibitor cocktail, and 2% Nonidet P40 substitute. Grinding took approximately 40-50 strokes, or until cuticle breakdown could be detected under a benchtop microscope. Following homogenization, protein was transferred into 1.5mL Eppendorf tubes and centrifuged at >13,000g for 20 minutes at 4°C, then returned to ice. The supernatant was transferred to a new tube and used to calculate protein concentration. For this step, BCA assay was used according to manufacturer’s instructions.

### Western blot

50µg of protein was used per sample. Upon transfer to a PVDF membrane, SYPRO Ruby was used to stain the membrane for total protein following manufacturer directions. After imaging, stained membranes were rehydrated in 100% methanol, briefly re-equilibrating in TBS + 1% Tween before adding blocking solution. 5% BSA in TBS-T was used for all blocking and antibody incubation steps. Anti-puromycin antibody clone 12D10 was used at a 1:2500 dilution, with secondary anti-mouse antibody at a 1:5000 dilution to detect puromycin. Each incubation was performed rocking at room temperature for 1-2 hours.

### Luciferase assay

Protocol adapted from (Olmedo et al., 2015). 300 worms per condition were treated with auxin until the early L4 stage. Worms were collected in S Basal and washed 3x to remove bacteria. In an opaque, flat-bottom 96-well plate, 100µL of S Basal was added to each well. 15 worms were pipetted into each well, including at least 5 replicate wells per condition. Once all worms were plated, a mix of 20g/L OP50 (wet weight), 200µM D-luciferin potassium salt, and 2.5μM 5-Ph-IAA or equivalent volume of DMSO for control was added to the relevant wells, taking care to fill all wells quickly for the same approximate start time for each condition. Worms treated with 20µM FCCP for one hour prior to plating and during the experiment were included as a positive control. Luminescence was detected using the Varioskan LUX plate reader, detecting for 1000ms once every 5 minutes for the duration of the experiment, approximately 1.5 hours total. Assistance with the plate reader was provided by Dr. Kim Lau and Paul Paroutis at the SickKids Imaging Facility.

### Lipid Quantification

Oil Red O (ORO) staining was performed on fixed worms as described in Escorcia et al., 2018. In brief, a working solution of 3 parts ORO stock solution (5mg/mL in 100% isopropanol) to 2 parts water was prepared, rocking overnight at room temperature protected from light. Remaining particulates were removed using a 0.2µm cellulose acetate syringe filter. 300 worms were collected from one medium NGM plate and washed 3x using PBS + 0.01% Triton X-100. Worms were dehydrated using 600µL 40% isopropanol, rocking at room temperature for 3 minutes. Supernatant was removed to 100µL, to which 600µL of ORO working solution was added. Samples were then protected from light and left to stain, rocking at room temperature, for 2 hours. Stained worms were washed once with 600µL PBS-T for 30 minutes to remove excess stain before mounting on slides. Imaging was performed using an AxioZoom.V16 microscope, fitted with an Axiocam 305 color camera. Assistance with the AxioZoom was provided by Dr. Kim Lau and Paul Paroutis at the SickKids Imaging Facility. Images were analyzed using ImageJ software. Whole worms were manually segmented, measuring signal in the green channel of the inverted image normalized to worm area.

### Temperature stress assay

50 synchronized worms were picked onto small NGM plates, with or without auxin according to the condition. Plates were incubated at 37°C for 3 hours and 30 minutes, removed from heat and allowed to recover at room temperature for 24 hours. Following this recovery period, worms were scored either dead or alive in response to physical prodding with a pick.

### Lifespan analysis

Worms were synchronized by bleaching in hypochlorite solution and transferred to auxin starting at 24 hours of age. Each plate was seeded with 100μL 10x concentrated OP50, to which approximately 50 worms were added. Every 2-3 days for the first 7 days worms were monitored for bacteria levels, transferring all conditions if any plate ran low on food. Worms were scored as dead when they became unresponsive to probing by pick. Deaths caused by bagging or exploded vulva were censored from the final data.

### Yolk quantification and intestinal width measurements

Worms expressing the AID2 system were crossed into the strain RT130 which carries the transgene pwIs23[vit-2::GFP]. Populations were synchronized by bleaching and approximately 300 worms were divided between auxin and control medium plates seeded with 10X concentrated OP50 at 24 hours of age. At each collection time, a subset of each population was slide mounted in 20mM tetramisole on 4% agarose pads and imaged at 10x in the eGFP channel and under brightfield using the Zeiss Axio Observer.Z1 microscope. Worms were outlined by hand in ImageJ, measuring total GFP signal and normalizing to body size. From the brightfield images, intestinal width was measured by drawing straight lines through the anterior intestine at three separate points and taking the average.

### Pharyngeal contraction rate

A population of approximately 100 synchronized worms was obtained by bleaching. Worms were transferred to small auxin or control plates at 24 hours of age. At each indicated age, worms were observed on their plates using a Leica MZ16 A stereomicroscope. After removing the plate lid, one minute was allowed for worms to acclimate. The number of pharyngeal contractions over the span of 30 seconds was counted manually, from which contractions per minute was estimated.

### Thrashing rate

Approximately 100 synchronized L1s were obtained by bleaching. Worms were transferred to small auxin or control plates at 24 hours of age. At each respective age, individual worms were collected by picking into the lid of a PCR strip tube containing 30μL M9 buffer with 0.01% Triton X-100 to prevent worms from sticking to the plastic. Once in solution, worms were allowed equilibrate to the liquid medium for 30 seconds before observation. Bending from the mid-body to one side was considered one body bend. Bends were counted by eye using a bench top microscope for a duration of 30 seconds, from which bends per minute was estimated.

### Statistical Analysis

All statistical tests were performed using Graphpad Prism 8. Student’s t-tests were unpaired, assuming Gaussian distribution. ANOVAs fit a full model and significance was calculated based on the appropriate multiple comparisons test. *p < 0.05, **p < 0.01, ***p < 0.001, ****p < 0.0001, ns = not significant.

## KEY RESOURCES TABLE

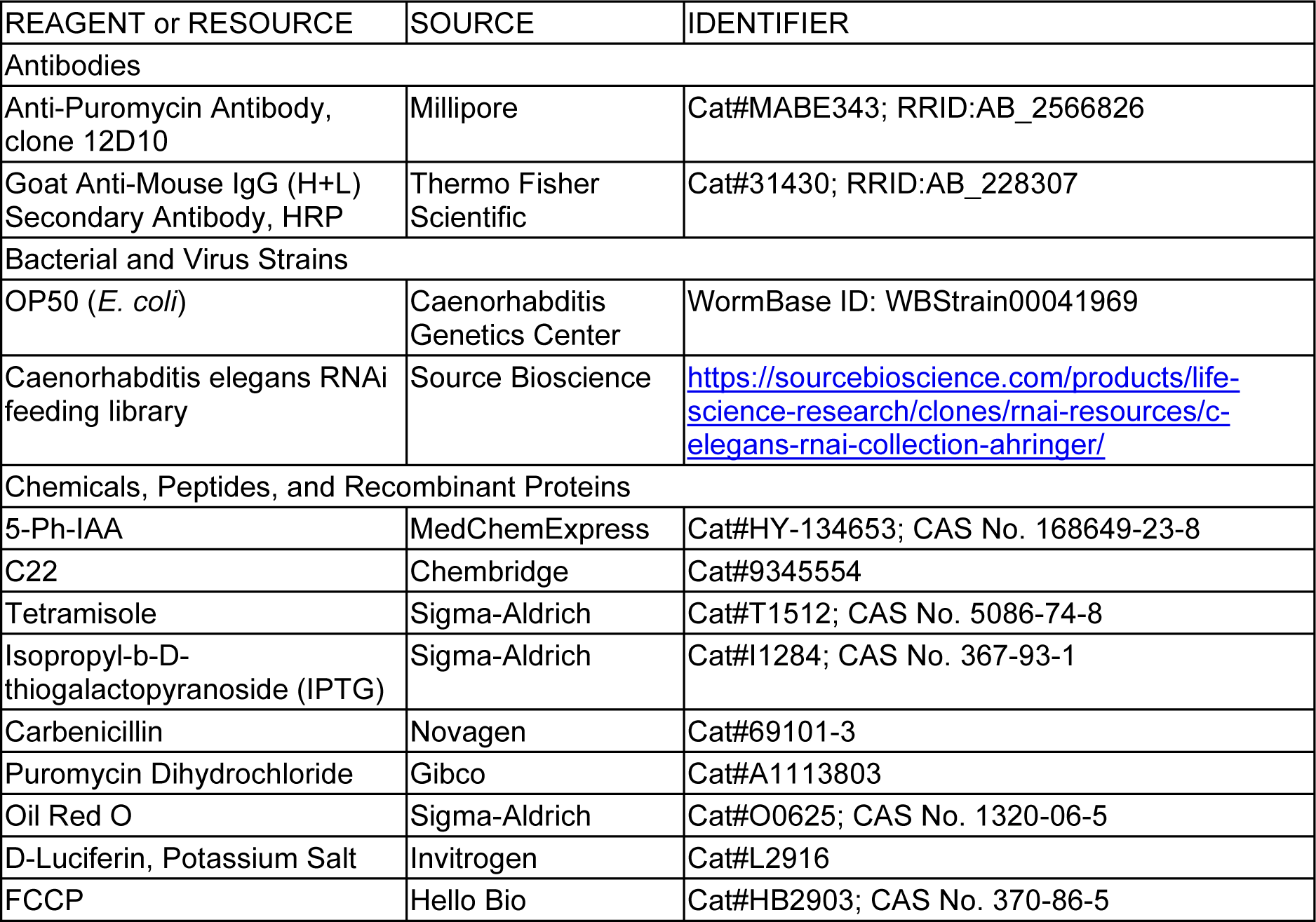
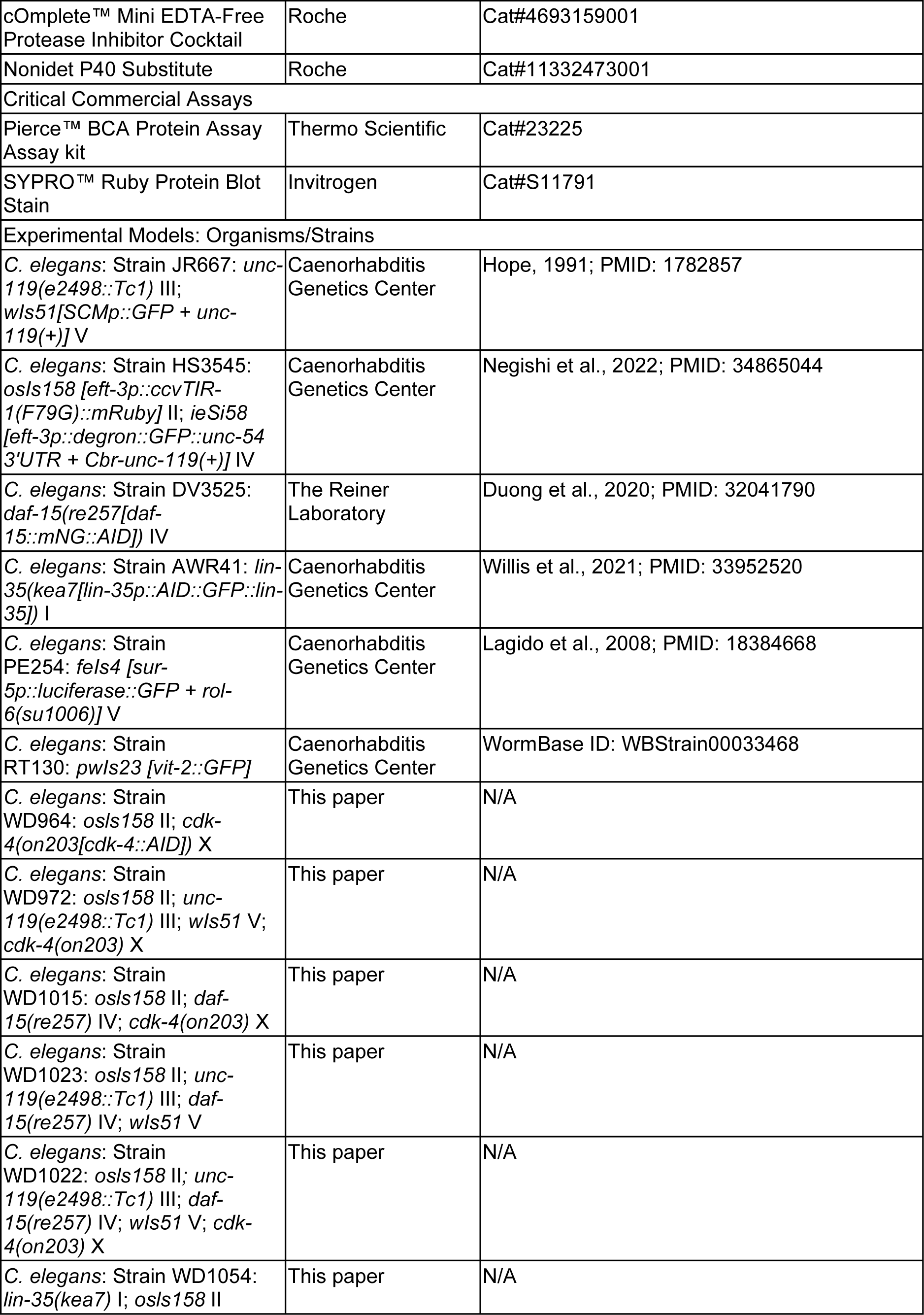
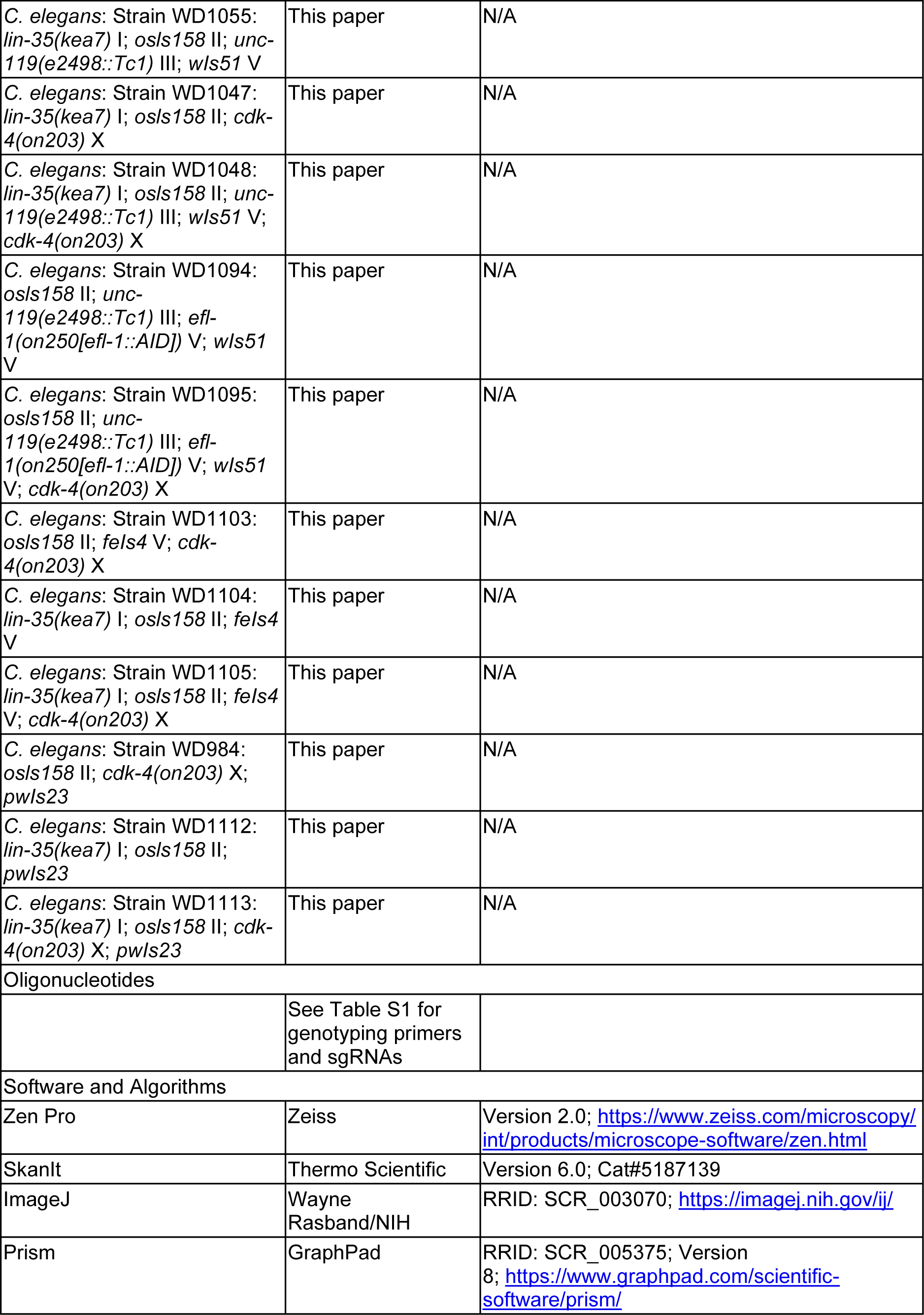
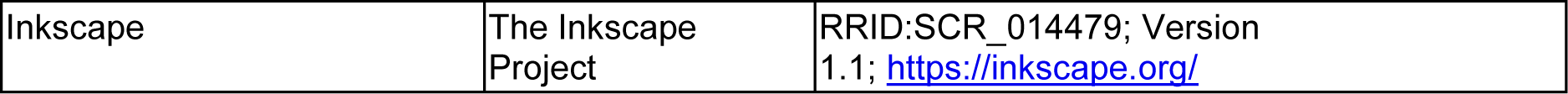

## Notes

### Competing Interest Statement

The authors have declared no competing interest.

### Summary of Updates

The only changes to this manuscript were in the discussion where we tightened up the language and edited some sentence structure issues.

