## Supplementary Information for "CDK-4 regulates nucleolar size and metabolism at the cost of late-life fitness in *C. elegans*"

**Supplementary Information:** CDK-4 regulates nucleolar size and anabolic metabolism as a fitness tradeoff in *C. elegans*.

Rachel Webster[1,3], Maria Quintana[2], Ran Kafri[1,3], W Brent Derry[2,3]

[1]Cell Biology Program, Peter Gilgan Centre for Research and Learning, The Hospital for Sick Children, Toronto, Canada.

[2]Developmental and Stem Cell Biology Program, Peter Gilgan Centre for Research and Learning, The Hospital for Sick Children, Toronto, Canada.

[3]Department of Molecular Genetics, University of Toronto, Canada.

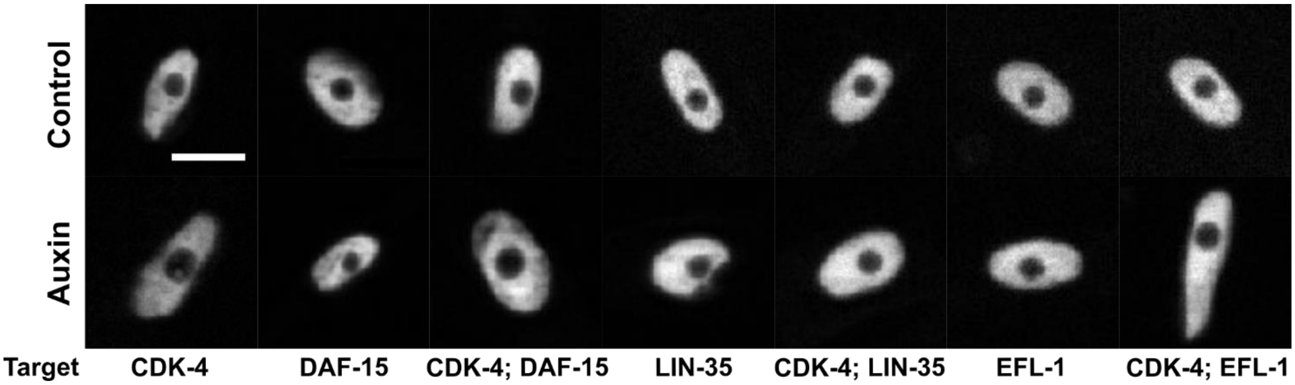


**Supplemental Figure 1: Representative images of nucleoli subject to indicated knockdown conditions.**

Control-treated examples are along the top row, with auxin-treated examples on the bottom for each of the labelled strains. Scale bar = 5μm.

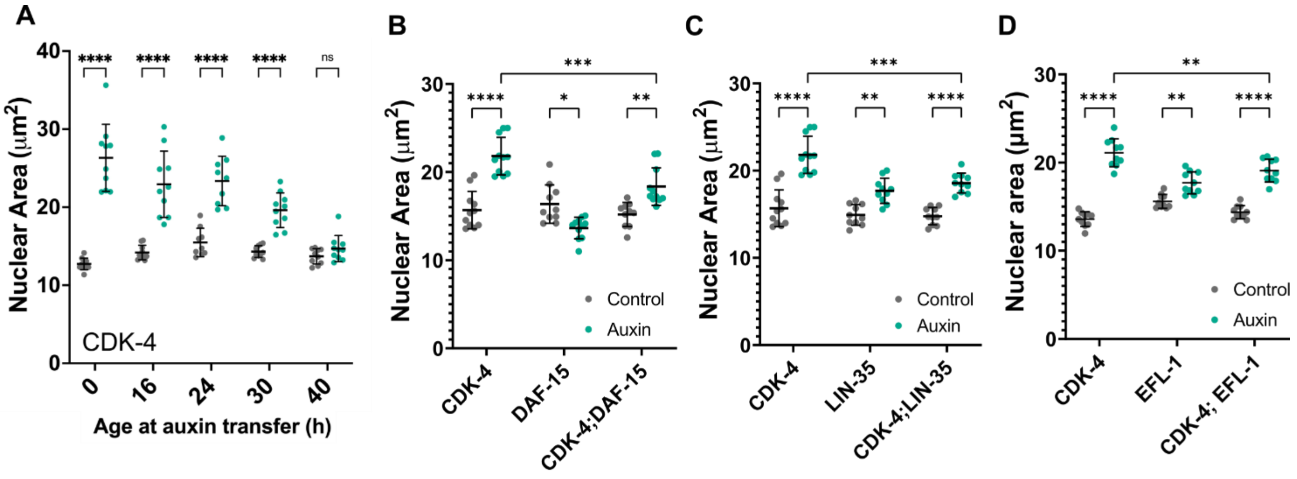


**Supplemental Figure 2: Nuclear size scales with nucleolar size.**

Nuclear size follows the same trends as seen with nucleolar size in **(A)** the time course experiment targeting CDK-4 at progressive starting points through development, and in all epistasis experiments examining interactions between **(B)** DAF-15, **(C)** LIN-35 and **(D)** EFL-1. Significance was calculated by two-way ANOVA with Šídák's multiple comparisons test in (A) and Tukey's multiple comparisons analysis in (B)-(D), with a minimum of 10 worms measured under all conditions.

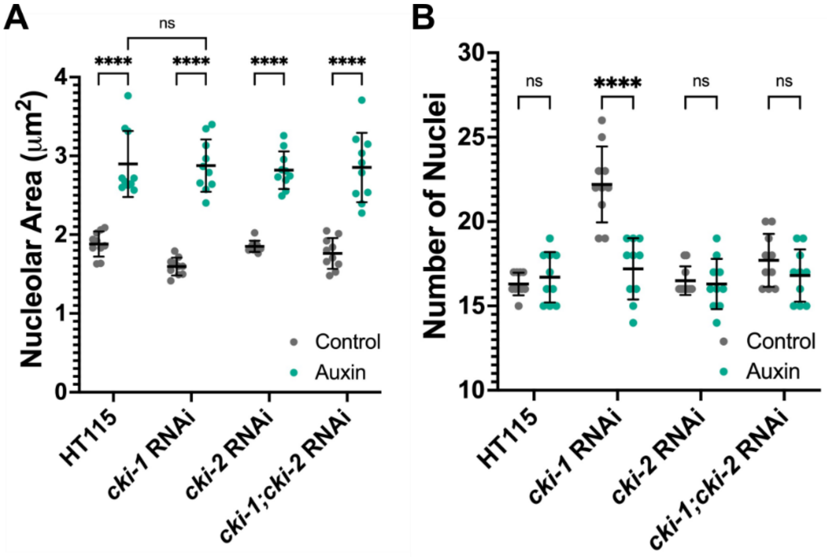


**Supplemental Figure 3: Interactions between CDK-4 and the CDK kinase inhibitors**

**(A)** Average nucleolar area and **(B)** number of nuclei of one seam syncytium, measured after two generations on the indicated RNAi, performed in *cdk-4*::degron worms. All groups were transferred to auxin or control RNAi plates at 24 hours of age. 10 worms across three biological replicates were measured per condition, with each data point representing the average measurement per worm. Significance was calculated using two-way ANOVA with Tukey's multiple comparisons analysis.

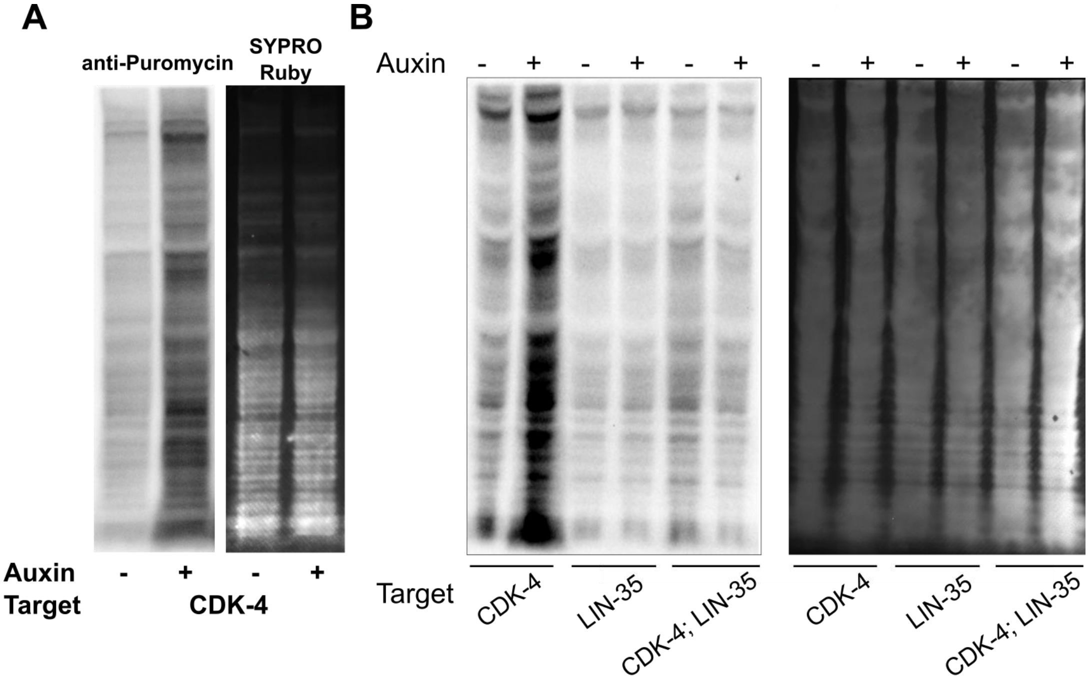


**Supplemental Figure 4: SYPRO Ruby loading controls for SUnSET assays.**

**(A)** Puromycin visualization by western blot (left) and corresponding SYRPO Ruby loading control (right) in control vs CDK-4 ablated populations. **(B)** Western blot probing for puromycin (left), with SYPRO Ruby loading control (right) for the indicated populations.

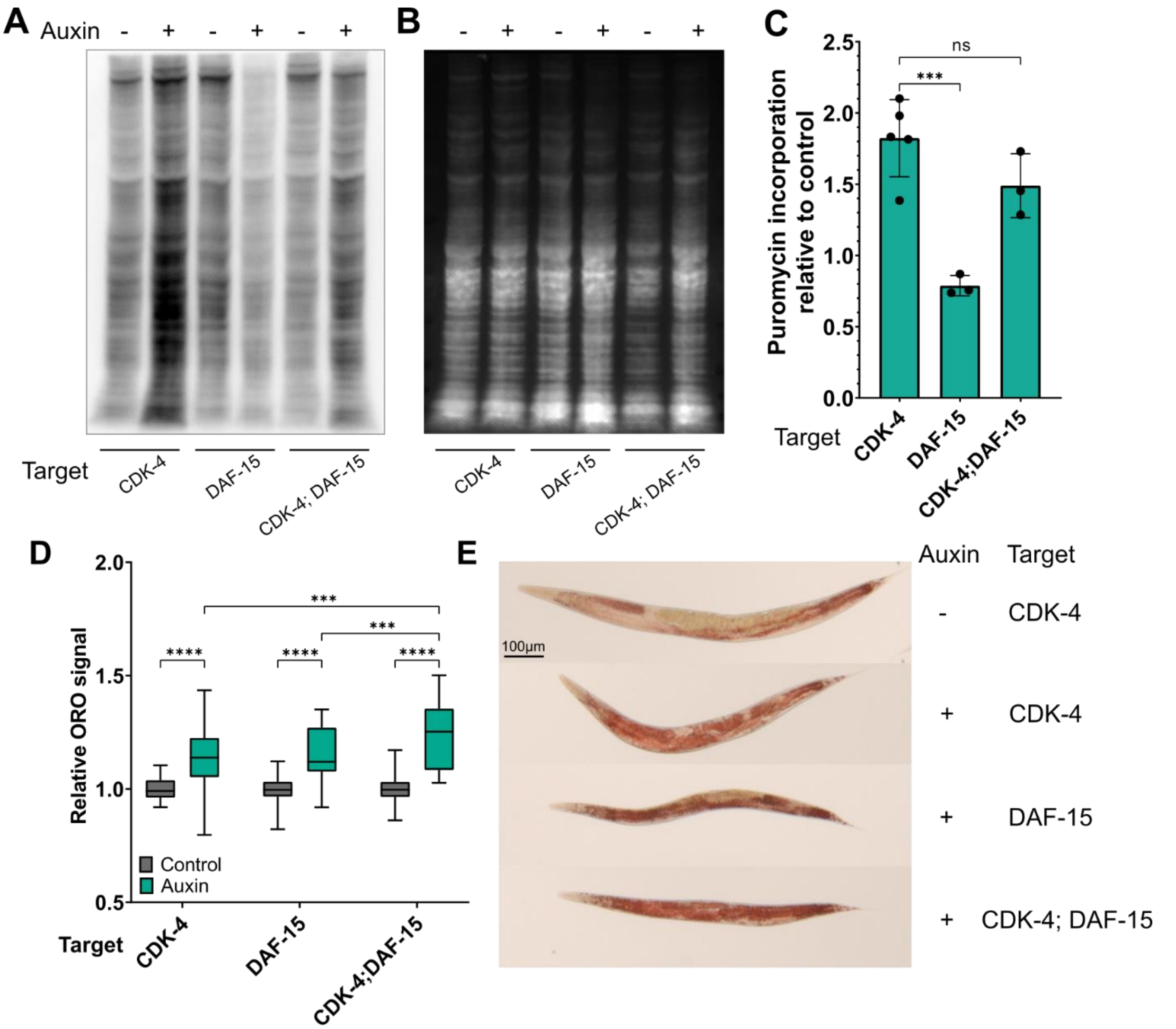


**Supplemental Figure 5: DAF-15 is not required for the metabolic phenotypes regulated by CDK-4.**

**(A)** Representative SUnSET assay western blot probing for puromycin using lysates collected from the indicated populations. **(B)** Corresponding SYPRO Ruby loading control. **(C)** Quantification of three biological replicates of the SUnSET assay. Data is represented as puromycin incorporation normalized to control values. Significance was calculated using one-way ANOVA with Holm-Šídák's multiple comparisons analysis. Bar height represents dataset mean, with error bars denoting standard deviation. **(D)** Quantification of Oil Red O staining, between 52-64 worms measured per condition taken from three biological replicates.

**Table S1: Oligonucleotides for generating and genotyping alleles.**

| Oligonucleotide | Sequence (5'-3') | Details |
|---|---|---|
| *cdk-4* gRNA | CCATCCGTTCTTGAAGCCTG | |
| *cdk-4* genotyping forward | CCAGTTGGCAAGGATTGGC | $T_m$ = 59.5°C,<br>507bp WT/639 KI |
| *cdk-4* genotyping reverse | CATTTCTGTCCGACTACCTCA | $T_m$ = 59.5 |
| *efl-1* gRNA | AGCACCGGAGACGAGTATCG | |
| *efl-1* genotyping forward | TCTCCGCCCACTTCTTCTGA | $T_m$ = 60.5°C<br>544bp WT/773 KI |
| *efl-1* genotyping reverse | AGGTTTTCTCCAGAGGCACA | $T_m$ = 58.4°C |
| *lin-35* genotyping forward | GAATTGTGGAACATCATTGCGC | $T_m$ = 60.1°C<br>560bp WT/1400 KI |
| *lin-35* genotyping reverse | GATTCGTTTTGCTGGTGGCTC | $T_m$ = 61.2°C |
| *daf-15* genotyping primer[1] | TCAGGAAGCAACTCCATCAATCG | $T_m$ = 62.9°C<br>900bp WT/600KI |
| *daf-15* genotyping primer[1] | TTGGAGACCATCGATGCTCC | $T_m$ = 60.5°C |
| *daf-15* genotyping primer[1] | CAAAACAAGACATGAAAATGGTGGCA | $T_m$ = 62.9°C |

[1]Primer sequences taken from Duong et al., 2020

**Table S4: Strains used in this study.**

| Strain name | Genotype | Source |
|---|---|---|
| JR667 | *unc-119* (e2498::Tc1) III; *wIs51*[SCMp::GFP + *unc-119*(+)] V | Hope *et al.,* 1991 PMID: 1782857 |
| HS3545 | *osIs158* [*eft-3*p::ccvTIR-1(F79G)::mRuby] II; *ieSi58* [*eft-3*p::degron::GFP:: *unc-54* 3'UTR + Cbr-*unc-119*(+)] IV | Negishi *et al.,* 2022 PMID: 34865044 |
| DV3525 | *daf-15*(*re257*[*daf-15*::mNG::AID]) IV | Duong *et al.,* 2020 PMID: 32041790 |
| AWR41 | *lin-35*(*kea7*[*lin-35*p::AID::GFP::*lin-35*]) I | Willis *et al.,* 2021 PMID: 33952520 |
| PE254 | *feIs4* [*sur-5*p::luciferase::GFP + *rol-6*(*su1006*)] V | Lagido *et al*., 2008 PMID: 18384668 |
| RT130 | *pwIs23* [*vit-2*::GFP] | CGC |
| WD964 | *cdk-4*(*on203*[*cdk-4*::AID]) X; *osls158* II | This study |
| WD1047 | *cdk-4*(*on203*) X; *lin-35*(*kea7*) I; *osls158* II | This study |
| WD1054 | *lin-35*(*kea7*) I; *osls158* II | This study |
| WD1015 | *daf-15*(*re257*) IV; *cdk-4*(*on203*) X; *osls158* II | This study |
| WD972 | *cdk-4*(*on203*) X; *osls158* II; *wIs51* V; *unc-119* (e2498::Tc1) III | This study |
| WD1023 | *daf-15*(*re257*) IV; *osls158* II; *wIs51* V; *unc-119* (e2498::Tc1) III | This study |
| WD1022 | *daf-15*(*re257*) IV; *cdk-4*(*on203*) X; *osls158* II; *wIs51* V; *unc-119*(e2498::Tc1) III | This study |
| WD1055 | *lin-35*(*kea7*) I; *osls158* II; *unc-119*(e2498::Tc1) III; *wIs51* V | This study |
| WD1048 | *lin-35*(*kea7*) I; *cdk-4*(*on203*) X; *osls158* II; *unc-119*(e2498::Tc1) III; *wIs51* V | This study |
| WD1094 | *efl-1*(*on250*[*efl-1*::AID]) V; *osls158* II; *wIs51* V; *unc-119* (e2498::Tc1) III | This study |

| | | |
|---|---|---|
| WD1095 | *efl-1*(*on250*) V; *cdk-4*(*on203*) X; *osls158* II; *wIs51* V; *unc-119* (e2498::Tc1) III | This study |
| WD1103 | *cdk-4*(*on203*) X; *osls158* II; *feIs4* V | This study |
| WD1104 | *lin-35*(*kea7*) I; *osls158* II; *feIs4* V | This study |
| WD1105 | *lin-35*(*kea7*) I; *cdk-4*(*on203*) X; *osls158* II; *feIs4* V | This study |
| WD984 | *cdk-4*(*on203*) X; *osls158* II; *pwIs23* | This study |
| WD1112 | *lin-35*(*kea7*) I; *osls158* II; *pwIs23* | This study |
| WD1113 | *lin-35*(*kea7*) I; *cdk-4*(*on203*) X; *osls158* II; *pwIs23* | This study |
